# A streamlined mass spectrometry-based proteomics workflow for large scale FFPE tissue analysis

**DOI:** 10.1101/779009

**Authors:** Fabian Coscia, Sophia Doll, Jacob Mathias Bech, Andreas Mund, Ernst Lengyel, Jan Lindebjerg, Gunvor Iben Madsen, José M. A. Moreira, Matthias Mann

## Abstract

Formalin fixation and paraffin-embedding (FFPE) is the most common method to preserve human tissue for clinical diagnosis and FFPE archives represent an invaluable resource for biomedical research. Proteins in FFPE material are stable over decades but their efficient extraction and streamlined analysis by mass spectrometry (MS)-based proteomics has so far proven challenging. Here, we describe an MS-based proteomic workflow for quantitative profiling of large FFPE tissue cohorts directly from pathology glass slides. We demonstrate broad applicability of the workflow to clinical pathology specimens and variable sample amounts, including less than 10,000 cancer cells isolated by laser-capture microdissection. Using state-of-the-art data dependent acquisition (DDA) and data independent (DIA) MS workflows, we consistently quantify a large part of the proteome in 100 min single-run analyses. In an adenoma cohort comprising more than 100 samples, total work up took less than a day. We observed a moderate trend towards lower protein identifications in long-term stored samples (>15 years) but clustering into distinct proteomic subtypes was independent of archival time. Our results underline the great promise of FFPE tissues for patient phenotyping using unbiased proteomics and prove the feasibility of analyzing large tissue cohorts in a robust, timely and streamlined manner.

## INTRODUCTION

Hospitals routinely archive biopsied tissue specimens that are collected for diagnostic purposes as formalin-fixed and paraffin-embedded (FFPE) tissue blocks. FFPE is an economical archival choice as tissues can be stored at great density at room temperature (RT) for years or decades while maintaining integrity for pathology analyses [1]. An estimated half a billion FFPE cancer tissues are stored to date and this number is rapidly rising [2], representing an invaluable resource for studying molecular mechanisms underlying diseases, testing potential biomarkers and discovering new ones. The long-term storage of FFPE specimens implies that they are often associated with a plethora of clinical data, including histology reports, treatment and patient outcomes. Such metadata are of particular importance for integration of retrospective clinical and molecular data to improve patient phenotyping and stratification. FFPE sample preparation involves denaturation, crosslinking and embedding in an inert matrix, which makes oligonucleotide based analysis difficult and often impossible [3,4]. Antibody-based histological methods are routinely performed but rely on the preservation of the relevant epitopes, which can also be very challenging and influence quantitation [5]. Because of these difficulties and the fact that mass spectrometry (MS)-based proteomics demands clean peptide sample to be introduced to the mass spectrometer, it came as a surprise when we and others showed that FFPE samples are indeed very well suited to this technology, including the analysis of post translational modifications (PTMs) [6,7]. Furthermore, carcinomas could be sampled to great depth, in excess of 10,000 expressed proteins [8]. Together these and other studies established that FFPE tissues show similar qualitative and quantitative proteomic properties as fresh frozen (FrFr) tissues, further highlighting the potential of FFPE tissue analysis for biomedical research [6,9,10].

Recent technological progress of MS-based proteomic methods now enable the unbiased and large scale investigation of proteomes at ever greater depth [11]. As a result, in-depth proteomic analysis of FFPE tissues in a clinical context has now become feasible. Using MS-based proteomics on archived FFPE tissues, we recently profiled more than 9,000 protein groups from metastatic ovarian cancer (OvCa) FFPE tissues and identified a novel prognostic disease biomarker, ‘CT45’, causally linked to patient long-term survival [12]. In another study, we stratified breast cancer patients into distinct proteomic subtypes based on newly discovered classification markers, some of which were not reflected at the mRNA level [13]. Using biobanked frozen tissue samples, studies from the Clinical Proteomic Tumor Analysis Consortium (CPTAC) highlighted the importance of integrating proteomic and phosphoproteomic data with genomic information to uncover functional consequences of genomic mutations and identify proteogenomic subtypes linked to patient outcome [14,15]. Despite these promising MS technology developments and improved protein extraction protocols from FFPE tissues, streamlined workflows for efficient processing of hundreds of tissue samples in a highly parallelized and robust manner are not yet available.

Protein analysis of FFPE tissue is challenging and has so far required complicated workflows to efficiently extract proteins from the formalin crosslinked tissue of interest. Hence, most proteomic studies including our own have so far focused on relatively small patient cohorts (less than 60 samples) and required large amounts of biological starting materials - often in the mg range. Typically, strong detergents such as sodium dodecyl sulfate (SDS) are used in lysis buffers, combined with boiling times up to a few hours to efficiently reverse formalin crosslinks, extract and solubilize proteins [16]. These methods, however, come with some caveats as detergents generally suppress trypsin activity and are incompatible with downstream LC-MS/MS analysis. This requires detergent removal steps that often cause sample loss. For instance, protein precipitation or filter devices can lead to significant protein losses, which also lowers reproducibility. Therefore, most detergent-based protocols are incompatible with ultra-low sample input, such as laser-capture micro-dissected (LCM) samples, which often range from only a few hundred to a few thousand cells. These sub-microgram protein amounts are easily lost in detergent clean-up steps and highlight the need for alternative protocols optimized for low sample amounts. Thus, a workflow enabling the processing of low as high-input samples, would be highly desirable. This is further illustrated from a pathologist’s perspective; depending on disease stage, anatomic location, and pharmacologic intervention or due to surgical circumstances, tissue availability can be highly variable and often limited. While pre-cancerous lesions can be comprised of only hundreds of cells, primary and metastatic tumors are typically available in larger quantities. Workflows that are not biased towards either end would therefore be of great clinical value for the investigating of disease mechanisms at the global proteome level.

Here, we report on our efforts to develop a robust, scalable and high-throughput workflow for quantitative proteomic profiling of patient derived FFPE tissue samples using different MS acquisition methods. We start with the observation that detergent-free protocols in principle offer an excellent alternative to existing workflows. Sample preparation is then carried out with MS compatible buffers in so-called ‘single-tube’ reactions to facilitate tissue lysis [17–19], protein extraction and tryptic digestion without additional clean-up or transfer steps. Organic solvents such as 2,2,2-trifluoroethanol (TFE) and acetonitrile (ACN) have overall good MS compatibility and yield high peptide and protein identifications. This is particular attractive for the analysis of low-input samples [18,20]. As a proof-of-concept, we applied our workflow to an adenoma cohort of more than 100 samples, thereby validating its applicability and value to reveal new insights into disease biology.

## RESULTS

### Proteomic sample preparation of archived biobank tissue in 96-well format

To be useful for large and diverse FFPE tissue cohorts, we reasoned that our proteomic workflow would need to be robust, streamlined and able to handle varying sample amounts. To this end, we optimized our recently described workflow for laser micro-dissected FFPE samples [21] to a 96-well format (Fig. 1A). We worked with macro- and micro-dissected FFPE sections on glass slides to test the applicability to varying FFPE tissue amounts. We here refer to ‘macro-dissections’ as razor blade scraped areas from FFPE sections, whereas by ‘micro-dissection’ we mean laser-capture microdissection (LCM) of tissue from glass membrane slides (Fig. 1B, Methods). We first macro-dissected similar tumor areas (~5 × 5 mm) by scraping from two consecutive 10 μm thick sections of the same high-grade serous ovarian cancer (OvCa) specimen. We then processed the collected tissues with our organic solvent (Methods) or SDS based workflow as described previously for in-depth proteomic profiling of FFPE biobank samples [12]. We noticed that long boiling (90 min) in the SDS buffer fully resolved FFPE tissue in contrast to the TFE-based lysis. After overnight tryptic digestion, however, the tissue was fully resolved with no noticeable undigested material. Following peptide clean-up, we found a 2.3-fold higher peptide yield using the TFE-based protocol (Fig. S1A), indicating both improved protein and peptide recovery [18,20] and protein loss during detergent clean-up steps. Next, we injected 0.5 μg of each peptide sample on a quadrupole Orbitrap mass spectrometer (Q Exactive HF-X) and analyzed them in 100 min single-run data dependent acquisition (DDA) mode (top15). Of the 5,041 identified protein groups, 92% were found with both protocols indicating that protein extraction was highly comparable despite the noticeable differences in tissue dissolving directly after boiling. The average proteome correlation was 0.95 (Pearson r) based on label-free quantification (MaxLFQ) values [22] (Fig. S1B). To assess differences in protein extraction, we calculated the proportion of proteins annotated to originate from different cellular compartments. This revealed nearly identical cellular compartment proportions for the two preparations of the same specimen (± 0.4%), and only minor differences (1-4%) compared to different FrFr OvCa tissues (Fig. 1C). We additionally analyzed differences in the extraction of chromatin bound proteins, which represent a particular challenge for formaldehyde fixed samples due to stable DNA-protein crosslinks that may hinder peptide identification. The relative protein abundance of chromatin and DNA binding proteins showed no significant difference for TFE and SDS processed samples (t-test *P* = 0.44 and 0.66, respectively), nor as compared to FrFr OvCa tissues (*P* = 0.33 for chromatin binding proteins, Fig. S1C). Together, these data demonstrate efficient protein extraction using our organic solvent-based workflow, including chromatin bound proteins.

**Fig. 1:**
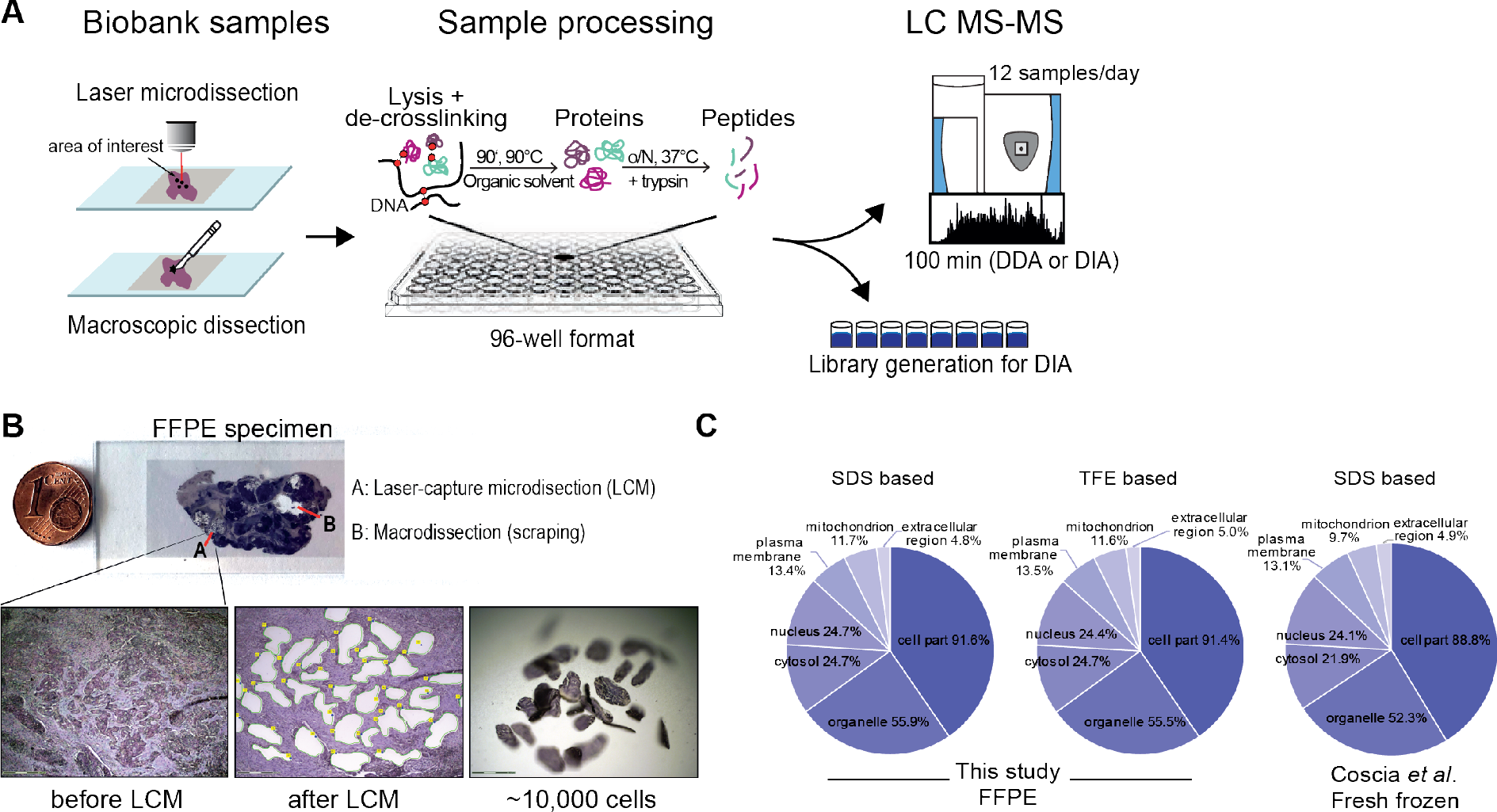
Overview of the mass spectrometry-based workflow for FFPE tissue analysis in 96 well format. **A)** Streamlined FFPE workflow. FFPE tissues are collected from glass slides either by scraping or laser capture microdissection (LCM), followed by in ‘single-tube’ sample processing and MS based proteomic analysis. **B)** FFPE tissue collection techniques. Minute sample amount can be either collected by LCM (A) or scraping (B) from a pathology glass slide. Images before and after LCM are shown in the first two lower panels. The third lower panel shows the collected LCM tissue regions, corresponding to an estimated number of 10,000 cancer cells (Methods), calculated based on the total volume of dissected tissue. **C)** Pie chart display the protein extraction efficiency from major cellular compartments (‘cytosol’, ‘nucleus’, ‘plasma membrane’ and ‘extracellular region’, Gene Ontology Cellular Component, GOCC) using two different lysis buffers containing SDS or TFE. Percentages reflect the number of proteins per compartment over all quantified proteins in the corresponding FFPE samples. The third pie chart shows the comparison to FrFr OvCa tissue proteome [29].

Formaldehyde crosslinking of FFPE tissue preserves tissue architecture, predominantly via stable methylene bridges between basic amino acid residues [23,24]. High proteome coverage of FFPE tissue therefore requires efficient formaldehyde de-crosslinking to reverse unwanted and variable chemical modifications that may obscure peptide identification. To assess differences in the reversal of formalin crosslinking between the two methods, we used pFind, an ‘open modification search’ algorithm [25]. This allowed us to screen for the most abundant protein modifications present in the analyzed FFPE tissues in an unbiased way. In line with previous reports [26,27], methionine oxidation and lysine methylation were the most abundant variable modifications compared to FrFr tissue (Fig. S1D). Importantly, we observed no apparent differences between SDS- and TFE-based sample lysis. However, we noticed up to 20% methionine oxidation of peptides with both protocols, indicating progressive protein oxidation over time [27]. Incomplete tryptic cleavage rates were similar between the methods with 24-26% and 1-4% of peptides displaying one or two missed cleavages, respectively (Fig. S1D).

Lysine methylation is a frequent protein modification in FFPE tissue, accounting for around 2-6 % of all identified peptides [26]. We also observed around 5% of these peptides in various FFPE tissues, even after prolonged de-crosslinking at high temperatures (90-100°C) and therefore reasoned that these ‘left-over’ peptides could be resistant to further de-crosslinking and thus indicate nearly complete crosslinking overall (about 95%). To ensure most efficient de-crosslinking, we opted for long heating times (90 min) in combination with ultrasound homogenization and high Tris concentrations (300 mM) in the lysis buffer [28] (Methods). Comparing the TFE- and SDS-based protocols at constant Tris concentration and boiling time resulted in similar lysine methylation rates (5.65 % vs. 4.89 %). We concluded that both protocols are equally efficient in terms of de-crosslinking. Of note, 74% of the lysine methylated peptides were exclusively identified in the methylated state. Using standard search engine settings, which do not include lysine methylation as variable modification, should therefore result in ~3-4% fewer peptide identifications (given no other unidentified modifications, or combination of those).

Among the lysine containing peptides that were identified in both methylated and unmethylated states, we observed that methylated sites were in median 70% less abundant (Fig. S1E). This observation was also true across diverse tissue types (Fig. S2C), indicating that differences in lysine methylation across samples should only marginally affect global proteome quantification.

Despite its beneficial chemical properties for improved protein and peptide extraction [18,20], TFE is a hazardous substance posing a higher safety risk compared to most other protein extraction methods. ACN is an alternative organic solvent, which shares similar physico-chemical properties while being less hazardous. Replacing TFE with ACN in our FFPE workflow resulted in similar peptide and protein yield, further reflected by high proteome correlations (Pearson r = 0.97) (Fig. S1F-H). Taken together, these results demonstrate that our sample processing protocol enables robust proteomic profiling of FFPE tissues with a detergent-free and ‘single-tube’ workflow.

### FFPE tissue workflow is broadly applicable to various tumor types

We next applied our workflow to FFPE sections from different tissue types (OvCa, glioma, colorectal adenoma and urachal carcinoma). Before collection, all tissue sections were subjected to microscopic inspection to identify neoplastic regions of interest that were subsequently macro-dissected or subjected to LCM (Fig. 1B, see Methods).

Excised tissue pieces were transferred into PCR tubes, allowing highly parallelized ‘single-tube’ sample preparation in a 96-well format (Figure 1A-B, S2A, Methods). MS analysis revealed excellent chromatographic reproducibility for micro- and macro-dissected samples with comparable total ion currents (TIC), independent of cancer origin or collection type (Figure S2B). We identified a similar number of peptides (on average 37,321) and protein groups (on average 4,933) for micro and macro-dissected samples in 100 min DDA single-run analysis, demonstrating the broad applicability of our workflow (Fig. 2A). To assess quantitative reproducibility, we compared the proteomes of five LCM FFPE tissues obtained from two advanced-stage high-grade serous OvCa (HGSOC) patients. We hypothesized that proteomic differences across patients would be larger than differences across anatomic sites from the same patient. Indeed, Pearson correlation coefficients were high for three biological replicates of the same patient (0.94-0.97), including primary (ovarian, OV) and metastatic (omental, OM) OvCa, and somewhat lower between patients (0.86 – 0.88) (Figure S2E). This result is consistent with our recent data demonstrating that the HGSOC tumor proteome is driven by patient-specific protein signatures independent of anatomic site [21].

**Fig. 2:**
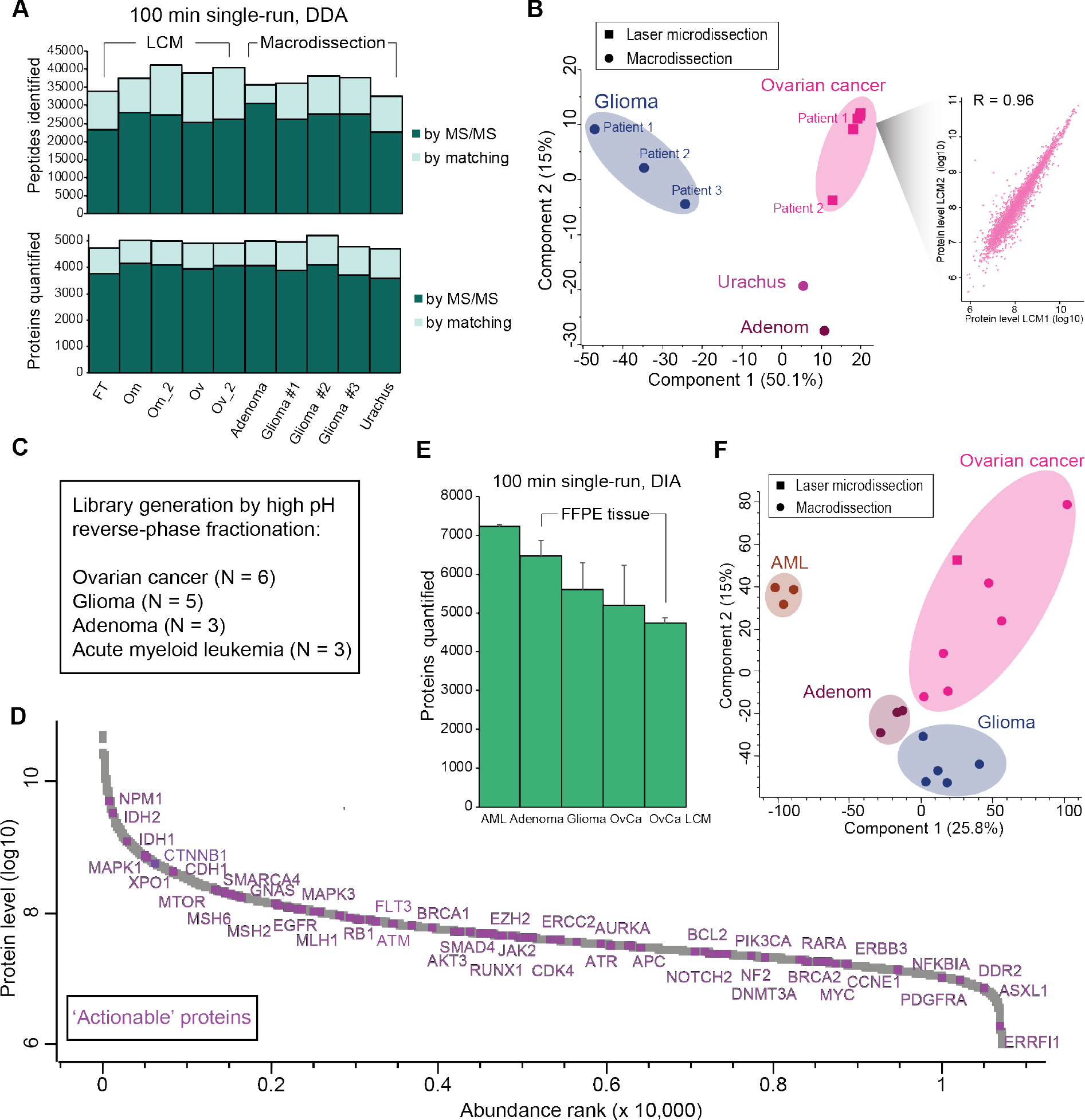
Evaluation of the FFPE tissue workflow in various tumor types. **A)**Bar plot of the total number of identified peptides (upper panel) and quantified proteins (lower panel) by MS/MS (dark green) and matching (light green) in invasive fallopian tube (FT) cancer, omental OvCa metastasis (OM), invasive OvCa (Ov), glioma, and urachal carcinoma (urachus). The first five tissues were collected by laser-capture microdissection (LCM) and the remaining five by macro-dissection. All samples were analyzed using 100 min single-shot DDA runs. **B)** Principal component analysis (PCA) of the ten tissue samples based on their proteomic expression profiles. The proteomes of glioma and OvCa samples are depicted by replicate number (patient 1-3 and collection 1-2, respectively). The first and second component segregate the samples and account for 50.1% and 15% of the variability, respectively. Ellipses encircle samples of same tissue origin. Reproducibility between two microdissected tissue samples from the same OvCa patient is depicted in the right panel (Pearson r = 0.96). **C)** Overview of the samples used for the generation of the spectral library by high pH reversed-phase fractionation. **D)** Dynamic range of the MS-signal of all identified proteins in the spectral library after high pH reversed-phase fractionation. Proteins belonging to the TARGET database, referred as actionable genes, are highlighted. **E)** Average number of quantified protein groups per tissue, including acute myeloid leukemia (AML), glioma and OvCa. Error bars represent standard deviations from minimum N = 3 replicates. **F)** PCA of tissue samples analyzed by 100 min DIA analysis. Ellipses indicate samples of same tissue origin.

Principal component analysis (PCA) of glioma, colorectal adenoma, OvCa and urachal carcinoma clearly grouped them according to their tissue of origin in the first and second components (Fig. 2B and S2D). The segregation was driven by known marker proteins such as the epithelial OvCa markers PAX8, MSLN, FOLR1 and MUC1. Brain-specific proteins like MBP, PLP1 and SLC1A2 were most abundant in glioma and the adenoma/urachus group showed high expression of the proteins AGR2, LGALS4, S100P and CDH17, which are known intestinal cell signaling and adhesion proteins (Fig. S2D). Quantitative completeness is an important aspect of proteomics and represents a particular analytical challenge in DDA strategies due to partially stochastic peptide sequencing. This is particularly true for single-run approaches that aim to quantify a large proportion of the cellular proteome without additional peptide pre-fractionation (which may not be appropriate for very low-input sample amounts or for reasons of limited total measurement time). To tackle this challenge, we first used the recently described ‘BoxCar’ scan mode, which allocates dynamic MS1 injection times based on precursor abundance [30]. Using BoxCar acquisition, the number of quantified protein groups increased on average by 7% in 100 min single-shot analysis resulting in 5,419 and 5,257 protein groups quantified in adenoma and glioma tissues, respectively (Fig. S2F).

With the recent increased scan speed of Orbitrap analyzers, this platform has become very attractive for DIA strategies [31]. In DIA, the entire sample complexity is in principle captured by cycling of the quadrupole selection window over the entire m/z range in pre-defined segments, thereby recording MS2 information irrespective of precursor intensity. While proteome coverage was previously a major bottleneck for DIA single-run workflows, more than 7,000 quantified protein groups have been reported from human cell lines in single 2 h measurements [32]. This prompted us to combine our streamlined tissue workflow with state-of-the-art DIA analysis based on the same 100 min LC gradient as in DDA analysis (Methods). Using high pH reversed-phase peptide fractionation [33], we generated a project specific spectral FFPE tissue library using our workflow encompassing 197,622 precursors, corresponding to 10,707 protein groups (Fig. 2C-D). The library covered a large number of known oncogenes and tumor suppressors with 108 members of the 135 <u>T</u>umor <u>A</u>lterations <u>R</u>elevant for <u>G</u>enomics-driven <u>T</u>herapy (TARGET) set reported as ‘actionable’ genes [34]. By definition, alteration of these genes or their expression levels can influence clinical decisions – making them particularly interesting in oncology.

Across tissue samples, we consistently quantified >5,000 protein groups and up to 7,268 protein groups for FrFr AML cells (Fig. 2E). The number of proteins matched to the library based only on single peptide hits was low with an average of 150 per sample. Of the 135 TARGET listed genes, almost half (44%, N = 59) were quantified on average in each sample. We conclude that our workflow provides a robust framework for proteomic profiling of FFPE tissue samples in 96-well format and is applicable to diverse tissue types, sample input amounts and MS-based proteomics strategies.

### FFPE tissue workflow shows high quantitative reproducibility

In addition to the robust analysis of laser micro-dissected samples, which showed high proteome correlations between biological tissue replicates (Fig. 2B and Fig. S2E), we assessed the reproducibility of our entire FFPE workflow for macro-dissected tissues. To this end, we collected tissue areas from glass slides, as routinely used in histopathology. Of note, LCM workflows instead require tissue mounting on specialized glass membrane slides (e.g. PEN or PET) that are generally not used in routine pathology. We focused on macro-dissections from colorectal adenomas to reduce biological complexity within and between sections as cancer lesions have high cellular heterogeneity at multiple levels [35]. Colorectal adenomas are benign precursor lesions to colorectal cancer.

They are genetically less complex and show chromosomal stability despite their large size [36]. To assess quantitative reproducibility, we repeated the entire workflow from sample collection and processing to MS measurement and data analysis. We mounted three consecutive tissue sections obtained from four different colorectal adenomas samples, all from the same patient, on glass slides. Based on histology, we collected the same tissue areas from all three adenoma sections using razor blade scraping. We reasoned that our biological inter-section comparison between consecutive 5 μM sections should be minor, allowing accurate estimation of the reproducibility of the entire workflow (Fig. 3A, left panel). Additionally, we collected three different areas from the same section to also assess intra-section proteome variability (Fig. 3A, right panel). Overall, we collected 19 different tissue samples that we processed on different days. In total, 6,110 protein groups were quantified in the single-shot 100 min DIA analyses, with an average of 5,274 per sample. Overall data completeness was 86% (Figure 3B). Single peptide hits comprised 345 protein groups on average, representing 6% of all quantified protein groups. To evaluate quantitative reproducibility, we calculated the coefficient of variation (CV) for all protein groups that were quantified in at least 70% of all samples (4,983 protein groups). CVs were calculated across similar regions from three consecutive sections of the same FFPE block (inter-section comparison) or from three different areas of the same tissue section (intra-section comparison, Fig. 3A and 3C). The inter-section variability was slightly lower than intra-section variability for all analyzed adenomas (average %CV of 18.9 and 20.1, respectively), whereas technical replicates of injections had a CV of 7.9% (Fig. 3A-D). Pearson correlations were correspondingly high with 0.91 – 0.95 for inter and intra-section replicates and 0.97 for technical replicates (Fig. S3A-B). As about 8% of total variation originates from the LC-MS measurement alone, we estimated that sample collection and processing should contribute no more than about 10% to the total observed variation, neglecting any true biological variation. Known adenoma or colorectal cancer markers such as E-Cadherin (CDH1), which is clinically used for patient diagnosis and stratification, was robustly quantified with our workflow with CVs below or close to 20% (Fig. 3D). Of note, even though EGFR was ~100-fold less abundant than COX-2 (MT-CO2), its variation was somewhat lower (17.7% vs 24.5%), indicating robust protein quantification even for lower abundant proteins. CV values across adenomas of the same patient (N=9) and across adenomas of different patients (N=9) showed higher proteome variations including known pathology markers, indicative of underlying differences in biology or pathology (Fig. 3E-F). Interestingly, ß-Catenin (CTNNB1) and CDH1 showed the lowest expression differences between sections within the same and different patients, highlighting their homogeneous expression patterns in adenomas as previously reported [37,38]. Conversely, EGFR and CDX2 were variably expressed between adenomas within and between patients, again in line with previous reports [39,40].

**Fig. 3:**
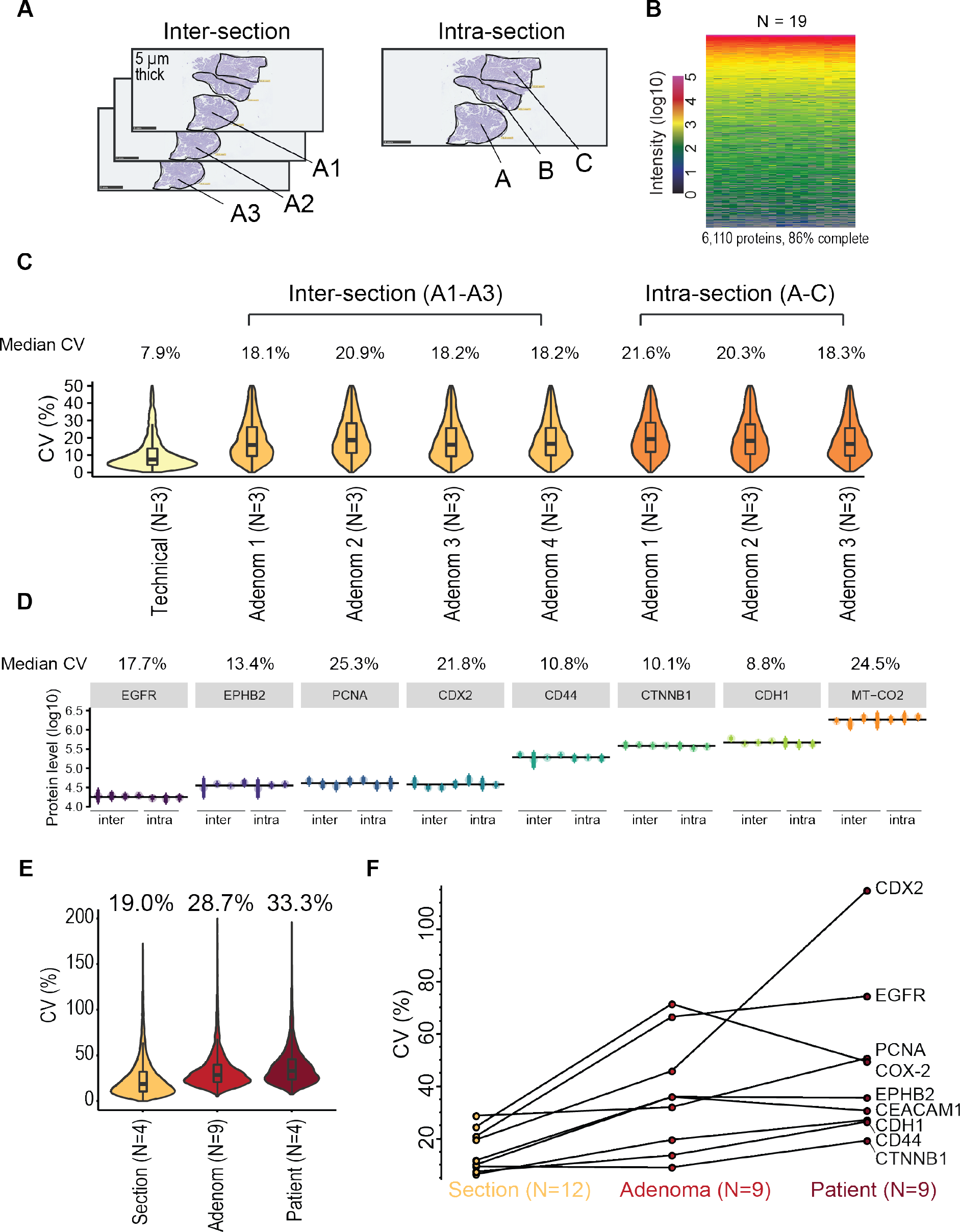
Evaluation of the reproducibility of the FFPE tissue workflow. **A)** Schematic representation of the H&E stained adenoma tissue sections used for overall workflow reproducibility assessment. Inter- and intra-section proteomic variability assessment is depicted in the left and right panels, respectively. Circles indicate the tissue regions manually collected by macro-dissection. **B)** Heatmap of protein abundance across inter- and intra-section tissue replicates from the same adenoma patient (N = 19 samples). In total, 6,110 protein groups are depicted, corresponding to 86% data completeness. **C)** Box plots and violin plots show the coefficient of variations (CVs in %) of protein quantification across injection replicates (N = 3), inter-section replicates (four sections of three replicates each) and intra-section replicates (three sections of three replicates each) based on 4,983 protein groups. For simplicity, values above 50% were excluded from the plots. CVs were calculated based on non-logarithmic values. **D)** Median CV values are plotted for eight known adenoma and colorectal cancer (CRC) markers across inter-(four values) and intra (three values)-section replicates. The average abundance for each protein across all sections is shown as a black line. Median CV values (%) for each protein across all sections are displayed above the plot. Note that CVs were calculated based on non-logarithmic values. **E)** CV values of protein quantification across adenoma sections (N = 4, same adenoma), adenoma tissues (N = 9, same patient) and patients (N = 9, different patients). **F)** CV values of nine quantified adenoma and CRC marker proteins across different sections, tissues and patients.

Together, these data demonstrate good workflow reproducibility and high proteome consistency within and across tissue sections of the same adenomas.

### Streamlined FFPE workflow applied to a cohort of 118 adenomas of different archival time

Having developed a scalable FFPE workflow applicable to diverse sample input amounts and cancer types, we next investigated if it would allow for streamlined processing of larger tissue cohorts within a single day and with minimal hands-on pipetting time. As a proof-of-concept, we analyzed an additional set of 118 FFPE adenoma tissues collected from 101 patients. After histopathological examination, samples were collected into 8-strip PCR tubes as before. All samples could indeed readily be processed in parallel within one day of sample preparation and subsequently scheduled for measurement within ten days using the single-shot 100 min DIA MS method (Methods). A total of 6,254 protein groups were quantified with a median of 5,147 protein groups per adenoma sample. We reached a data completeness of 92% for 5,230 quantified protein groups and 81% for all 6,254 protein groups (Fig. 4A and S3C).

**Fig. 4:**
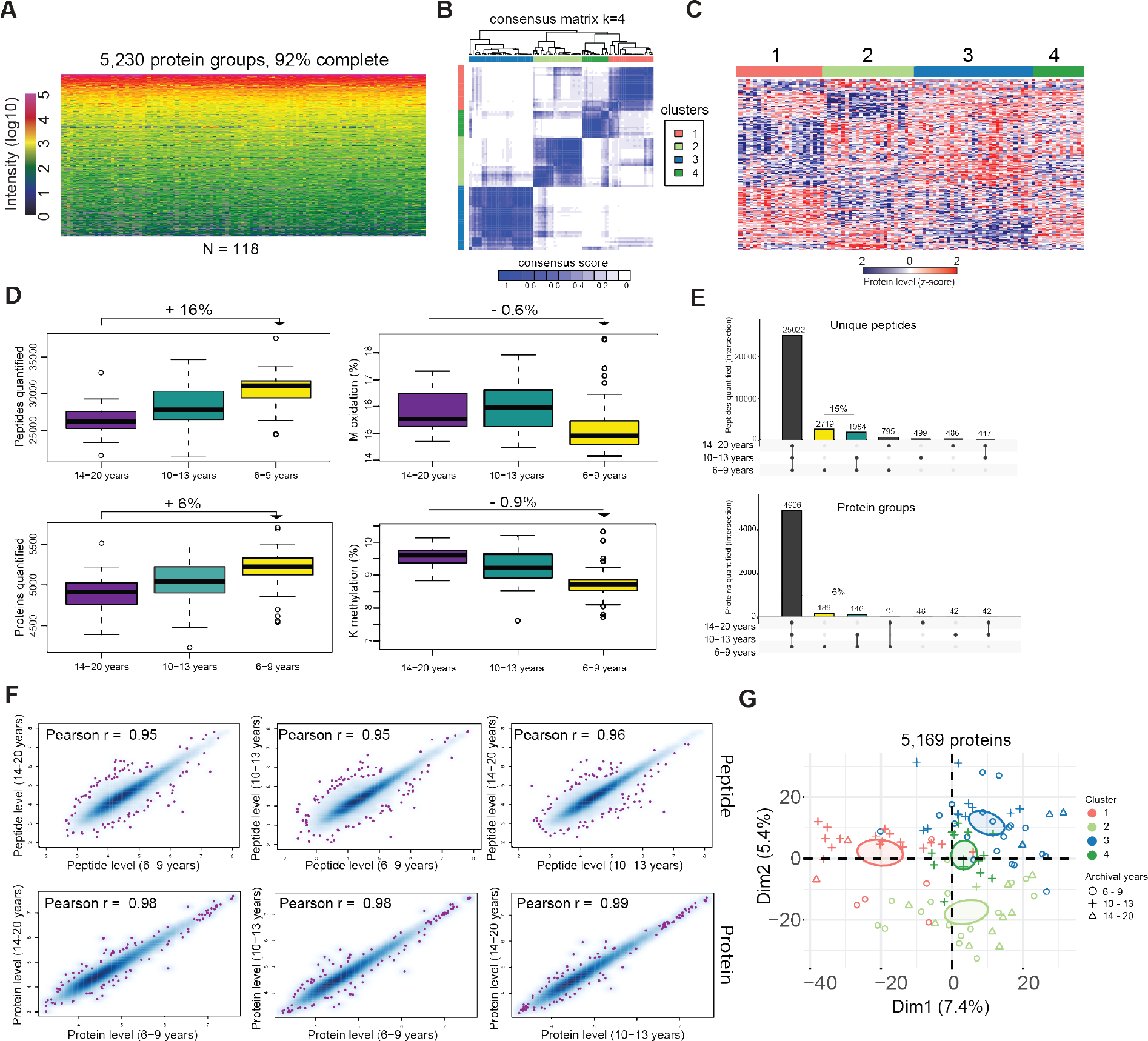
Streamlined proteomic analysis of 118 adenoma tissues of varying archival time. **A)** Heat map of protein abundance across 118 adenoma tissues encompassing 5,230 protein groups. **B)** Consensus clustering of the proteomic expression data determined by Pearson correlation as distance metric, where k represents the total number of clusters. Consensus scores are indicated using a color scale from white (samples never cluster together) to blue (samples always cluster together). **C)** Heat map of z-scored protein abundances of the 416 differentially expressed proteins (ANOVA, FDR < 0.01, s_*0*_ = 0.5) after unsupervised hierarchical clustering reveals four distinct adenoma clusters. Up- and down-regulated proteins are represented in red and blue, respectively. Euclidean distance was used for row-wise clustering. **D)** Box plots displaying the total number of quantified peptides and proteins, as well as lysine methylation and methionine oxidation in FFPE tissues archived with different storage times, ranging from 6 to 20 years. Samples were grouped according to archival years into three groups: 14-20 years (N=14), 10-13 years (N=41) and 6-9 years (N=43). For each group, a minimum of 50% quantified values was required. **E)** UpSet plots showing intersections of peptide (upper panel) and proteins (lower panel) quantification across archival time groups. **F)** Scatterplot of pairwise proteomic comparisons between archival groups on peptide (upper panels) and protein level (lower panels). **G)** PCA of 101 colorectal adenoma samples based on their proteomic expression profiles, encompassing 5,169 proteins. The first and second dimension segregate the samples and account for 7.4% and 5.4% of the total variability, respectively. The color code corresponds to the four adenoma subclusters identified in panel 4B whereas symbol types indicate the three archival groups. Point concentration ellipses are shown for each cluster with a 95% confidence interval.

This relatively large number of adenoma tissue proteomes prompted us to investigate the molecular differences across adenomas and the impact of archival time on the proteome. The latter is of particular relevance for the design of large discovery-based studies to avoid potential sampling biases. To identify the optimal number of proteomic subgroups in our dataset, we applied a consensus clustering approach [41] based on the 1,000 most variably expressed protein groups across adenomas as measured by median absolute deviation. Three samples were excluded from the analysis due to low protein identifications. This resulted in four major proteome clusters of similar size, which showed unique protein expression profiles (Fig. 4B, C and Fig. S3D-F Fig. 4C, ANOVA FDR < 0.05). We observed some overlap across the four clusters, indicative of shared biological features (Fig. 4C). For instance, proteins related to mitochondria and cellular respiration were significantly higher in clusters 1, 3 and 4 versus cluster 2, whereas cell adhesion and immune system related proteins were higher in clusters 2, 3, and 4 versus cluster 1(Fig. S3F).

Next, we investigated the impact of storage time on the four identified proteomic clusters. Tissues had been collected and archived between the years 1998 and 2008 with a median archival time of 11 years and a maximum of 20 years. Samples were grouped by archival time into 14-20 years (N=14), 10-13 years (N=41) and 6-9 years (N=43). We first compared the total number of quantified peptides across archival time groups, which revealed a somewhat higher number of quantified peptides for the short-term group versus the mid-term (−10%) and long-term group (−16%) (Fig. 4D), independent of sample amount as reflected in the total ion current (Fig. S3H). This trend was also apparent when we plotted the number of peptides against archival time as a continuous variable (Fig. S3I). At the protein level, however, this trend was less pronounced with losses of only −4% and −6%, respectively. To investigate whether the lower identification rates are related to increased methionine oxidation and lysine methylation over time (the two most abundant variable peptide modifications in FFPE tissue), we included lysine methylation as a variable modification in the spectral library search and re-analyzed the DIA data. This revealed a general trend towards a higher peptide modification rate in the long-term storage samples, indicative of progressive protein modification over time [27], however, at 1% this difference was much too small to explain the reduction of 16% in peptide identifications in the oldest samples (Fig. 4D). A previous study reported lower identification rates in tissue samples stored more than 20 years, and linked it to compromised retrieval of low abundant proteins [42].

We likewise found that the peptides exclusively quantified in the short-term storage group (2,719 peptides, 8% of all peptides), were significantly lower abundant than the peptides shared across groups (Fig. 4E, Fig. S3G). A similar abundance trend was observed at the protein level but to a much lower extent (−6%, 189 protein groups unique to the short-term group and 146 to the short-term and mid-term group compared to the long-term group (Fig. 4E, Fig. S3G). Filtering for the 3,000 most abundant protein groups in our dataset (calculated by median protein level across all samples) resulted in only 1% less protein quantifications in the long-term storage group (Fig. S3J). The impact on global protein quantification was also minor as judged by the high proteome correlations between different archival age groups (Pearson correlations 0.95-0.99) (Fig. 4F). Our data are consistent with previous proteomic studies showing that protein quantification is generally not perturbed even after 30 years of storage [6,10,42]. Given these findings, we analyzed how different archival times were represented in the four adenoma subclusters, which are depicted in Fig. 4B. Reassuringly, biological variability between samples was more dominant than variability caused by archival time, and as a result archival groups were sub-divided into the four separate clusters (Fig. 4G). This further illustrates the general applicability of FFPE tissues for proteomics-centered disease phenotyping and patient stratification.

## DISCUSSION

FFPE is the most common method to store tissues in pathology due to its long-storage ability, simplicity and economy. Hundreds of millions of FFPE samples exist worldwide in tissue biobanks, awaiting analysis. These immense archives represent an invaluable resource and opportunity to study molecular mechanisms of the diverse diseases for which tissue samples are taken. MS-based proteomics has increasingly been used over the last years to investigate FFPE samples at the proteome level but this typically required tedious workflows, lacked reproducibility, throughput and also analytical depth. This prompted us to develop a streamlined and robust MS-based workflow for the proteomic investigation of many FFPE samples.

With our new workflow in hand, we demonstrated the practicality of analyzing relatively large clinical FFPE cohorts from pathology glass slides with good proteomic depth in single-run analysis using state-of-the-art DDA and DIA acquisition methods. Based on a single-tube workflow in 96-well format and using MS compatible protein extraction buffers, we demonstrated high proteome similarity to detergent based sample preparation and achieve a proteomic depth of 5,000 – 6,500 protein groups per single sample. As tissue amounts are often highly variable in histopathology, it was critical to make our workflow applicable also to low sample amounts. To this end, we showed that laser-microdissection samples, encompassing less than 10,000 cells, can be analyzed in a reproducible fashion. We also demonstrated that FFPE tumor types of different entities can be reproducibly processed with our workflow as required in pathology practice. We retrieve known biological differences between OvCa, glioma and urachal carcinoma, validating the overall quality of our workflow. We also cover a large proportion of known ‘actionable’ proteins after high-pH reversed phase fractionation, suggesting that the proteomic analysis of FFPE tissues will be clinically relevant in the future.

Our short sample preparation time of less than one day, followed by prompt MS measurement and data analysis, highlights the promise of our FFPE workflow in clinical pathology practice in the future, where fast sample analysis for diagnosis and target identification in patients is very important. It is pertinent that our workflow can be well integrated with routine tissue preparation in pathology where direct tissue collection from glass slides of H&E stained sections offers several important advantages. Separate de-paraffinization, which usually comes at the cost of throughput and sample preparation time, is not needed, as it is already part of standardized H&E staining procedures. Furthermore, the availability of image data, collected routinely in histopathology for patient diagnosis, can be integrated to guide sample collection and to avoid contamination from non-diseased tissue areas, which is often a major limitation for bulk tissue analysis. This is illustrated in our analysis of adenoma tissues where we found a high degree of proteomic similarity between overlapping tissue areas across sections of the same FFPE block. In this tissue type, we further investigated the impact of archival time on several proteome quality parameters including peptide and protein quantifications, as well as peptide modifications. While the literature reports conflicting results on the impact of archival time on the number of identified peptides and protein groups, we noticed a 16% drop in peptide identifications in long-term storage samples (14-20 years), compared to 6-9 years old samples. However, one 15 year old sample showed a similar number of peptide identifications compared to the average of the 6-9 years old samples (Fig. S3I), suggesting that good tissue processing and storage could alleviate this effect [27,43]. In any case, the drop in peptide quantification only translated to 6% less quantified protein groups, indicating a general ‘buffer’ effect on protein levels that sustains global proteome integrity of old samples. This highlights the advantage of shotgun proteomic approaches that typically integrate information from multiple peptides for protein quantification. Our data further demonstrate that the quantitative protein information was not perturbed by storage time as we observed high global proteome correlations between storage groups. This suggests that previously published conflicting results might have arisen from differences in proteome coverage due to the workflows employed. Filtering our dataset for the most abundant proteins revealed a drop of only 1% of protein quantifications between the different storage groups. While additional investigation will be required to fully address the impact of archival time on the proteome, however, we found that biological differences across adenoma tissues drives the segregation of samples independently of archival times. Our analysis revealed previously unknown adenoma clusters, reflecting potential new subtypes that could lead to a novel classification. This speculation, however, will require a more thorough molecular analysis to validate these preliminary data.

In the future, our proteomic FFPE workflow can be extended with phosphoproteomics for FFPE samples, which will require streamlining of the analyses at the post-translational modification (PTM) level as well as even higher sensitivity. This will be important because information at the phosphorylation level can be crucial to understand disease biology, for instance as it reflects kinase activity driving tumor development. Furthermore, in combination with genomic information, multi-omic data could further improve current molecular diagnosis [44]. To this end, promising developments of high-throughput enrichment methods, using robotic assistance, have been developed and are becoming more practical. For instance, automated phospho-peptide enrichment can be achieved in only one hour [45], representing a promising future avenue for FFPE tissues analysis. Finally, single-cell technologies are becoming increasingly important to profile genetic, epigenetic, spatial, proteomic and lineage information of individual cells [46]. We believe that a variation of such single cell approaches could also be developed for FFPE analysis. For all these reasons streamlined, robust and scalable workflows for FFPE tissue analysis, such as the one introduced here, will be of immense value.

In conclusion, we here demonstrated a high-throughput and reproducible proteomic workflow, which creates the possibility for researchers to analyze large clinical FFPE tissue cohorts with wide ranging sample amounts in a robust, timely and streamlined manner. We hope that will help to trigger the proteomic analysis of thousands of valuable FFPE tissues. This information can then be combined with other ‘omic’ data with the ultimate goal of uncovering tissue biomarkers for better patient classification, diagnosis or treatment.

## Supporting information

Sample Preparation Protocol

## Acknowledgements

We thank all members of the department of Proteomics and Signal Transduction at the Max Planck Institute of Biochemistry and the Clinical Proteomics Group at the The Novo Nordisk Foundation Center for Protein Research for support and fruitful discussions. In particular, we thank Jakob Bader and Arend Koch (Charité Berlin) for assistance with glioma tissue analysis, Philipp Nuhn, Maximilian C. Kriegmair and Stefan Porubsky (Mannheim medical center) for providing urachal FFPE samples, Annette Bartels for assistance with colorectal adenoma tissue collection and preparation, Ashok Kumar Jayavelu and for the assistance with AML collection and analysis, and Philipp Geyer for sample preparation input. This work was supported by the Max Planck Society for the Advancement of Science and by the Novo Nordisk Foundation (grant agreement NNF14CC0001 and NNF15CC0001), European Union’s Horizon 2020 research and innovation program under grant agreement 686547 (MSmed Project) and the University of Chicago Cancer Center Support Grant P30CA014599.

## Supplementary data and Methods

**Supplementary figure 1:**
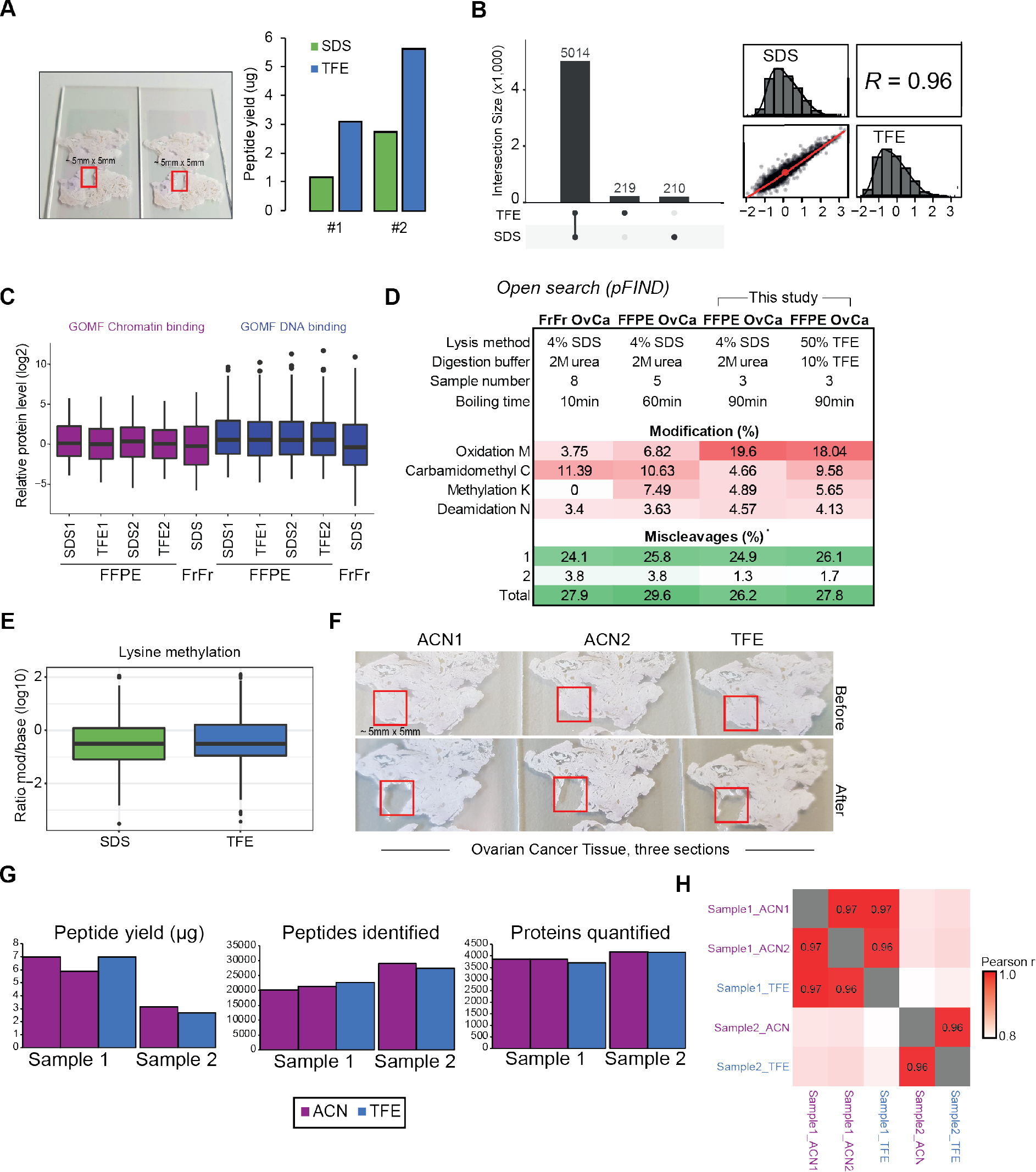
Mass spectrometry-based proteomic workflow for FFPE tissues. **A)** Comparison of SDS and TFE-based protocols. Left panel: Example pictures of OvCa tissue section replicates of the same tumor (10 μM thick) used to compare TFE and SDS based protocols. Red boxes indicate the collected areas by scraping (approximately 5 mm × 5 mm). Right panel: Total peptide yield (μg) after clean-up by StageTips measured on a Nanodrop instrument. **B)** Quantitative comparison of SDS and TFE-based protocols. Left panel: UpSet plot showing intersections of protein quantification across the two protocols. Right panel: pairwise proteome comparison of tissue replicates processed with the SDS or TFE-based protocol. Protein levels are median centered and log10 transformed with a Pearson correlation (R) of 0.96. **C)** Comparison of relative protein abundance between SDS and TFE-based protocols. Box plots show relative protein abundance (sample median normalized) of ‘Chromatin binding’ or ‘DNA binding’ (Gene Ontology Molecular Function, GOMF) proteins. OvCa proteomes obtained from FrFr tissue [29] is plotted for relative comparison to non-formalin fixed tissue. **D)** Open modification search (pFIND) results of FFPE tissues. The table summarizes results the most abundant peptide modifications and tryptic miscleavage rates of OvCa tissues (panel A) processed with the SDS or TFE-based protocol. Previous OvCa studies are included as comparison. FrFr OvCa [29], FFPE OvCa [12]. **E)** Comparison of lysine methylation rates between SDS- and TFE-based protocols. The ratios of lysine methylated peptide intensities to unmodified base peptide intensities are shown as boxplot. **F)** Comparison of ACN and TFE-based protocols. Pictures of three OvCa tissue section replicates of the same tumor (10 μM thick) used to compare ACN and TFE-based protocols. Red boxes indicate areas before and after collection by scraping (approximately 5 mm × 5 mm). **G)** Proteomic results for two OvCa tissue samples processed in replicates with the ACN or TFE-based protocol. Sample 1 corresponds to sections shown in panel F. **H)** Proteome correlation map (Pearson r) for two OvCa tissue samples processed in replicates with the ACN or TFE-based protocol.

**Supplementary figure 2:**
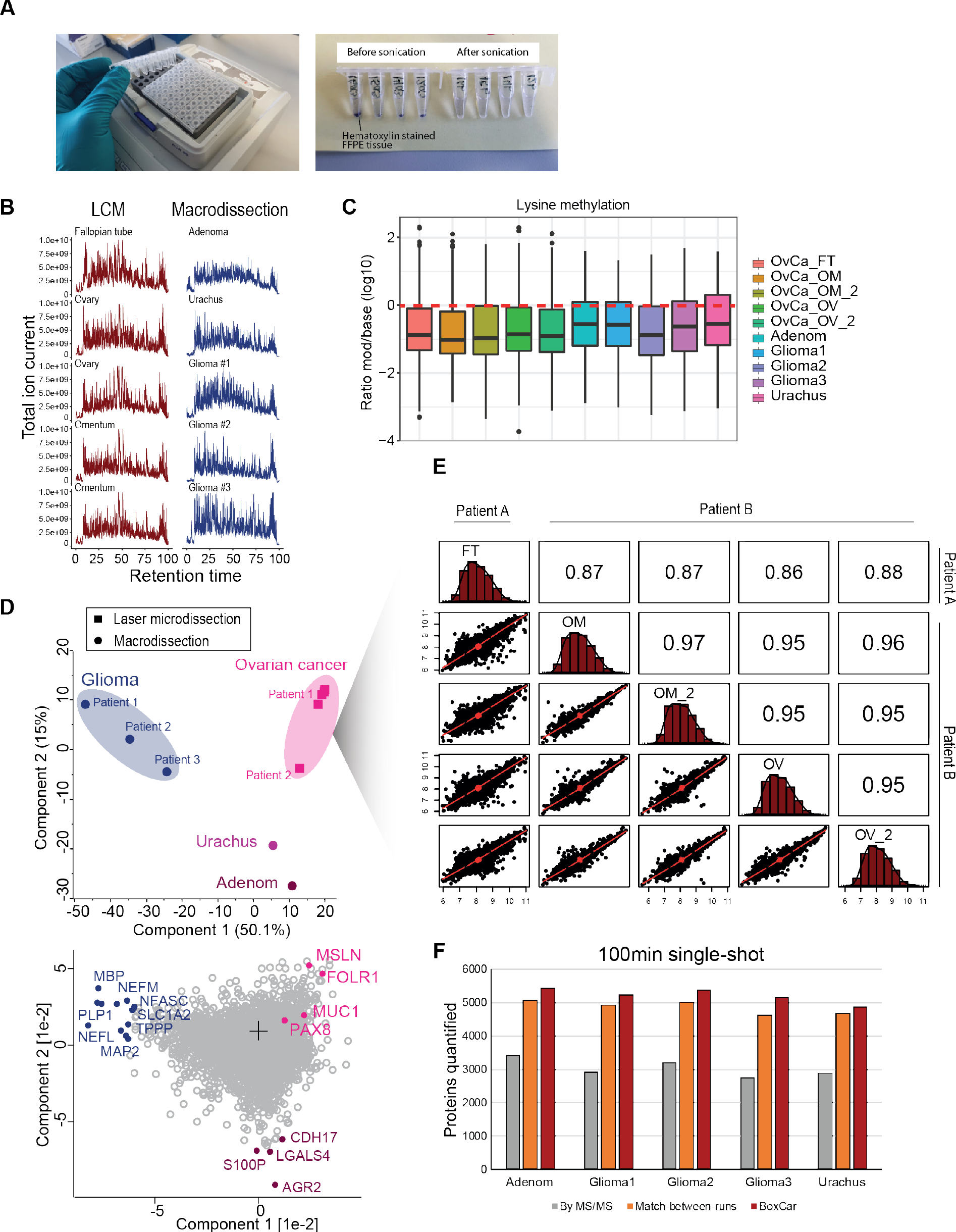
FFPE tissue workflow is broadly applicable across tissue types. **A)** Streamlined FFPE tissue workflow in 96-well format. The left panel shows how 96 samples are processed with our tissue workflow. The right panel depicts PCR tubes with H&E stained tissue in lysis buffer before and after sonication. **B)** Total ion current of the mass spectrometric analysis of tissue samples collected by LCM or macro-dissection. **C)** Lysine methylation rates across ten FFPE tissue samples. The ratios of lysine methylated peptide intensities to unmodified base peptide intensities are shown as boxplot. **D)** Principal component analysis (PCA) of ten FFPE tissue based on their proteomic expression profiles. Upper panel: The first and second component segregate the different FFPE tissues and account for 50% and 15% of the variability, respectively. Samples collected by LCM and macro-dissection are depicted as rectangles and dots, respectively. Ellipses indicate samples of same tissue origin. Proteins driving the segregation between the different FFPE tissues are depicted in the lower panel. Several known protein markers for the corresponding tissues are highlighted. **E)** Proteome correlation matrix of five laser microdissected OvCa tissues obtained from two patients with high-grade serous OvCa. Depicted values are Pearson correlations. FT: invasive fallopian tube cancer, OM: omental OvCa metastasis, Ov: invasive OvCa, GBM: glioma, Urachus: urachal carcinoma. **F)** Total number of quantified protein groups for macro-dissected tissues in 100 min single-shot DDA analysis. Grey bar plots represent the protein groups only identified by MS/MS, orange bar plots the protein groups identified by matching (‘match-between-runs’), and red bar plots the protein groups identified by ‘BoxCar’ acquisition.

**Supplementary figure 3:**
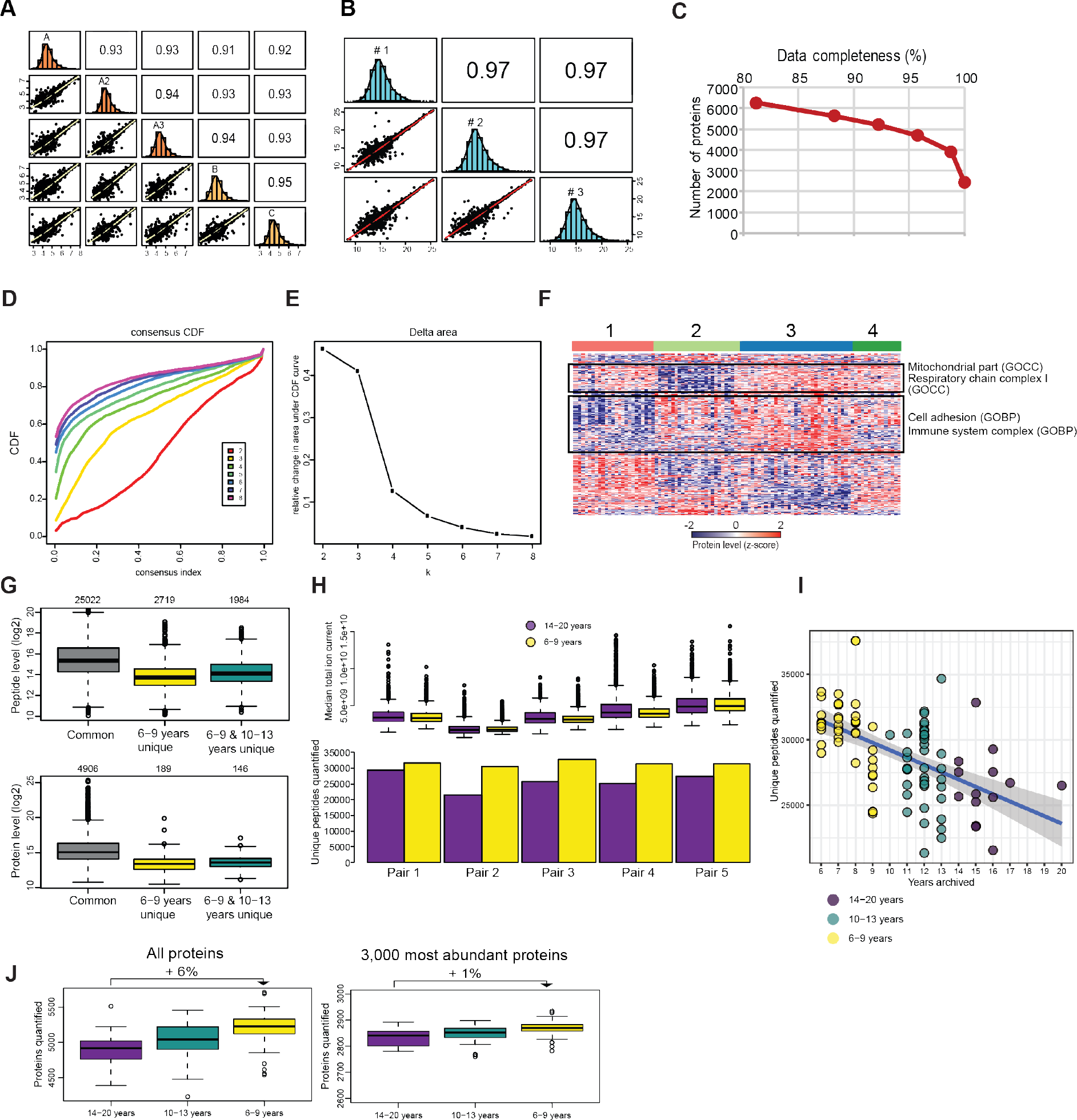
Reproducible and streamlined proteomic analysis of adenoma tissues of varying archival time. **A)** Proteome correlation matrix of inter-(A1-A3) and intra-section (A-C) replicates of FFPE adenoma tissues. Depicted values are Pearson correlations. **B)** Proteome correlation matrix of three injection replicates (#1-3) of the same adenoma sample, with corresponding Pearson correlations. **C)** Total number of quantified protein groups in the adenoma tissue cohort (N=118) in relation to data completeness. **D)** Consensus Cumulative Distribution Function (CDF) plot. The cumulative distribution functions of the consensus matrix are plotted for each number of clusters, k, (indicated by colors), estimated by a histogram of 100 bins. The plot allows the determination of an optimal number of clusters, k, where the CDF reaches an approximate maximum with consensus and cluster confidence at a maximum. **E)** Delta area plot showing the relative change in area under the CDF curve comparing k and k − 1. For k = 2, no k −1 is present, explaining why the total area under the curve rather than the relative increase is plotted. This plot enables the determination of the relative increase in consensus and k at which there is no or little increase. **F)** Co-regulated pathways across the four adenoma clusters. Heat map of z-scored protein abundances of the 416 differentially expressed proteins (ANOVA, FDR < 0.01, s_*0*_ = 0.5) after unsupervised hierarchical clustering, related to Fig. 4C. Up- and down-regulated proteins are represented in red and blue, respectively. Significantly enriched pathways (Benjamini-Hochberg FDR < 0.05, GOCC and GPBP terms) are shown exemplarily for co-expressed protein clusters across the four patient groups. **G)** Intensity of commonly and uniquely quantified peptides (upper panel) and protein groups (lower panel) in the three archival time groups. **H)** Effect of archival time on the number of quantified peptides. Upper panel: Box plots showing total ion currents (TIC) of the samples used for pairwise comparison. Samples of different archival groups (6-9 years and 14-20 years) were matched by their median TIC (calculated for the main peptide elution window 10-80 min in the 100 min gradient). Lower panel: Bar plots showing the number of quantified peptides for each sample. Sample pairs matched by TIC are depicted next to each other. **I)** Scatter plot showing the number of quantified peptides *versus* archival years. A linear model-based trend line is shown including 95% confidence intervals (grey area). **J)** Total number of quantified proteins in FFPE tissues archived with different storage times. All proteins that were quantified in each group with at least 50% valid values are shown in the left panel, whereas the 3,000 most abundant proteins in the dataset (calculated by median protein level across all 98 samples) are plotted in the right panel.

## METHODS

### Sample collection

All samples used in this study were human FFPE tissues and collected with informed consent (University of Chicago Institutional Review Board-approved protocol [13372] for OvCa, Charité University Hospital Berlin for glioma [EA2/101/08], University Hospitals Jena for AML [Vote 3477-06/12], University Medical Centre Mannheim for urachal carcinoma [2015-540-MA], Health Research Ethics Committee of the Capital Region of Denmark for colorectal adenomas [H-16022392]) and in accordance with the Declaration of Helsinki.

A detailed sample collection and preparation protocol is provided in the supplementary material. Briefly, FFPE tissues were sectioned using a microtome (5 or 10-μm sections). Tissue slices were mounted on regular or Leica PEN-membrane Membrane Slides (2 μm) pathology glass slides for macro-dissection or LCM, respectively. Slides were deparaffinized with xylene and rehydrated through graded alcohols and water. Sections were stained with Mayer’s hematoxylin (Sigma) and dehydrated through graded alcohols and xylene. Areas of interest were determined via microscopic inspection by pathologists.

For macro-dissections, an area of roughly 5 mm × 5 mm was collected by scraping with a razor blade (5 μM or 10 μM thick section) and transferred into 0.2 ml 24-well PCR tubes (Thermo Scientific, cat. no. AB-0624). In contrast, micro-dissected tissues were dissected with a Zeiss PALM laser microdissection system and collected into adhesive caps. A total area of approximately 2.5 × 10E6 μm^2^ was collected by LCM. This area corresponds to about 10,000 cells, as derived from dissected area × slide thickness / average mammalian cell volume of 2,500 μm3, BNID 100434.

For AML cell collection, newly diagnosed untreated de novo AML patient cells were obtained in accordance with the Declaration of Helsinki. The peripheral blood samples were enriched for mononuclear cells using density-gradient centrifugation for 30 mins at 400 × g using BIOCOLL separating solution (Density 1.077, Biochrom, Berlin Germany). The blast-enriched PBMCs were washed in 1X PBS once and cryopreserved at −150C.

### Sample preparation for MS analysis

Both macro and micro-dissected tissue pieces were transferred into 0.2 ml 24-well PCR tubes (Thermo Scientific, cat. no. AB-0624) containing 100 μl lysis buffer (50% 2,2,2-trifluoroethanol [TFE], 300 mM Tris/HCl pH8).

PCR tubes were tightly closed with 8-cap strips (Thermo Scientific, cat. no. AB-0784). Samples were further sonicated (15 cycles at high intensity using a Bioruptor, Diagenode) and boiled (90°C, 90min) in a ThermoMixer (Eppendorf) using a 96-well adapter. Next, samples were briefly centrifuged to collect condensate and sonicated a second time. After short centrifugation, proteins were reduced with 5 mM 1,4-dithiothreitol (DTT) for 20 min (1,500 rpm) followed by alkylation with 25 mM 2-chloroacetamide (CAA) for another 20 min (1,500 rpm). Lysates were then vacuum concentrated for 45min at 60 °C (until max. 20 μl remained). Protein digestion was performed overnight at 37°C and 1500 rpm by addition of 80 μl digestion buffer (10% TFE in ddH20 with trypsin and LysC at an enzyme/protein ratio of 1:50). On the following day, samples were shortly centrifuged and digestion was stopped by adding trifluoroacetic acid (TFA) to a final concentration of 1%.

Peptides were then desalted and purified with styrenedivinylbenzene-reversed phase sulfonate (SDB-RPS) StageTips [47] as follows. Peptides were directly loaded on 2 layers of SDB-RPS and centrifuged for 5 min at 500xg. After one wash with 200 μl 1% (vol/vol) TFA in isopropanol, and one wash with 200 μl of 0.2% TFA in H2O, peptides were eluted with 50ul elution buffer (80% ACN and 1% ammonia) and vacuum-dried completely. Dried peptides were finally reconstituted in 2% ACN and 0.1% TFA in water and kept at −20°C until MS analysis.

The matching library was prepared using the same protocol and was comprised of FFPE tissue of OvCa, adenoma and glioma, as well as AML and fractionated using the high pH reversed-phase fractionator as previously described (Kulak et al. MCP). Briefly, we fractionated a total of about 30 μg of peptides that was automatically concatenated into 8 or 24 fractions using a rotating valve that switches the elution flow every 90 sec.

### LC-MS/MS analysis

Nanoflow LC–MS/MS analysis of tryptic peptides was conducted on a quadrupole Orbitrap mass spectrometer (Q Exactive HF-X, Thermo Fisher Scientific, Rockford, IL, USA) [48] coupled to an EASY nLC 1200 ultra-high-pressure system (Thermo Fisher Scientific) via a nano-electrospray ion source. 300 ng of peptides were loaded on a 50-cm HPLC-column (75-μm inner diameter, New Objective, USA; in-house packed using ReproSil-Pur C18-AQ 1.9-μm silica beads; Dr Maisch GmbH, Germany).

Peptides were separated using a linear gradient from 2 to 20% B in 55 min and stepped up to 40% in 40 min followed by a 5 min wash at 98% B at 350 nl per min where solvent A was 0.1% formic acid in water and solvent B was 80% ACN and 0.1% formic acid in water. The total duration of the run was 100 min. Column temperature was kept at 60 °C by an in-house-developed oven.

For DDA analysis, the mass spectrometer was operated in ‘top‐15’ data‐dependent mode, collecting MS spectra in the Orbitrap mass analyzer (60,000 resolution, 300–1,650 m/z range) with an automatic gain control (AGC) target of 3E6 and a maximum ion injection time of 25 ms. The most intense ions from the full scan were isolated with an isolation width of 1.4 m/z. Following higher‐energy collisional dissociation (HCD) with a normalized collision energy (NCE) of 27, MS/MS spectra were collected in the Orbitrap (15,000 resolution) with an AGC target of 1E5 and a maximum ion injection time of 28 ms. Precursor dynamic exclusion was enabled with a duration of 30 s. The DIA method consisted of one MS1 scan (350 or 300 to 1,650m/z, resolution 60,000 or 120,000, maximum injection time 60 ms, AGC target 3E6) and 32 segments at varying isolation windows from 14,4 m/z to 562,8 m/z (resolution 30,000, maximum injection time 54 ms, AGC target 3E6). Stepped normalized collision energy was 25, 27.5 and 30. The default charge state for MS2 was set to 2.

### MS data analysis

DIA raw files were analyzed with Spectronaut Pulsar X software (Biognosys, version 12.0.20491.17 under default settings for targeted DIA analysis with ‘mutated’ as decoy method. We used a projects specific spectral library encompassing 197,622 precursors, corresponding to 10,707 protein groups. Data export was filtered by ‘No Decoy’ and ‘Quantification Data Filtering’ for peptide and protein quantifications. For adenoma tissue analysis (Fig. 3 and 4), we used an adenoma tissue specific library, encompassing 7,725 protein groups (77,275 precursors).

DDA raw files were processed in the MaxQuant environment [49] (version 1.5.0.38). The integrated Andromeda search engine [50] was used for peptide and protein identification at an FDR of less than 1%. The human UniProtKB database (October 2017) was used as forward database and the automatically generated reverse database for the decoy search. ‘Trypsin’ was set as the enzyme specificity. Search criteria included carbamidomethylation of cysteine as a fixed modification, oxidation of methionine, acetyl (protein N-terminus) and lysine methylation (as stated) as variable modifications. We required a minimum of 7 amino acids for peptide identification. Proteins that could not be discriminated by unique peptides were assigned to the same protein group. Label-free protein quantification was performed using the MaxLFQ [22] algorithm and ‘match-between-runs’ was enabled. We used a minimum ratio count of one peptide and filtered out proteins with only one razor or unique peptide. Proteins, which were found as reverse hits or only identified by site-modification, were filtered out.

### Statistical analysis

All statistical and bioinformatic analyses were performed using Perseus [51] or the R framework (https://www.r-project.org/). Missing values were imputed based on a normal distribution (width = 0.3; downshift = 1.8). Consensus clustering was performed based on the ConsensusClusterPlus R library [41]. The 1,000 most variably expressed protein groups (calculated by median absolute deviation) were used for consensus clustering. The number of clusters, k, was varied from 2 to 8 with 1,000 resamplings. Hierarchical clustering was used based on Pearson correlations as distance metric. The Consensus Cumulative Distribution Function (CDF) plot and Delta Area plot were used to assess the optimal number of clusters.

## Author contributions

The study was conceived by F.C, S.D and M.M. Proteomic sample preparation, analysis and interpretation were performed by F.C. and S.D, under supervision of M.M. Experiments were designed by F.C, S.D. OvCa tissues were collected and selected for the study by E.L. and microdissected by F.C. Adenoma tissues were collected and selected for the study by J.L. and G.I.M. and prepared for proteomic sample preparation by J.B., and processed by F.C. and A.M. Figures were prepared by F.C. and S.D. The paper was written by F.C, S.D and M.M. The paper was edited by F.C., S.D., J.B., A.M., E.L., J.M., and M.M. All authors reviewed and provided feedback on the manuscript.

