## Supplementary material for "A streamlined mass spectrometry-based proteomics workflow for large scale FFPE tissue analysis": Sample Preparation Protocol

---

Protocol version 1.0

Submitted to bioRxiv on Sept 23[rd], 2019

**<u>Disclaimer</u>:**
**This protocol includes hazardous chemicals and any use of it is strictly at your own risk**

**(!) Read safety information**

### Detailed sample preparation protocol

#### REAGENTS

• (!) 2,2,2-Trifluoroethanol (TFE; Sigma-Aldrich, cat. no. 96924-250ML-F)
• (!) 1,4-dithiothreitol (DTT; Roche, cat. no. 10197777001)
• (!) 2-Chloroacetamide (CAA; Sigma-Aldrich, cat. no. C0267)
• Tris-HCl (hydroxymethylaminomethane hydrochloride)
• Endoproteinase Lys-C (Wako Chemicals, cat. no. 129-02541)
• Proteomics grade modified trypsin (Sigma-Aldrich, cat. no. T6567).
• (!) Acetonitrile (ACN; Fisher Scientific, cat. no. A955-4)
• (!) Isopropanol (ISO; Fisher Scientific, cat. no. A461-1)
• (!) Trifluoroacetic acid (TFA; Merck, cat. no. 8082600100)
• (!) Ammonia solution, 25% (NH4OH; Merck, cat. no. 5330030050)

#### EQUIPMENT

• Eppendorf ThermoMixer C (Eppendorf, cat. no. 5382000015) with 96-well adapter for PCR tubes.
• 8-channel 200 µl pipette and 8-channel 10 µl pipette.
• 0.2 ml 24-well PCR plate (Thermo Scientific, cat. no. AB-0624).
• Flat 8 Cap Strips (Thermo Scientific, cat. no. AB-0784).
• In-house made 96-well 'swimming' adapter for Bioruptor based sonication of PCR tubes. Note, plastic inlets from 200 µl tip boxes can be used for this purpose.
• Solid-phase extraction disks for SDB-RPS StageTips: Empore SDB-RPS (Sigma, cat. no. 66886-U). We use in-house made StageTips [1].
• StageTip Centrifuge STC-V2 (https://www.sonation.com/en/products/STZentrifuge/index.html). Alternatively, if many samples are prepared, we recommend a 3D printed device capable of holding 96 StageTips. A 3D printer employing fused deposition modelling such as the Zortax M200 or Ultimaker 3 instruments enable the rapid (~12 hour) and inexpensive fabrication of these devices. If such a device is not available, a pipette-tip box may serve as a suitable StageTip holder.
• Evaporative concentrator: Eppendorf Vacuum Concentrator Plus with 96-well plate rotor.

#### REAGENT SETUP

**Stock solutions:**

1M Tris/HCl pH8 in ddH2O
500 mM 2-Chloroacetamide (CAA) in ddH2O
100 mM 1,4-dithiothreitol (DTT) in ddH2O
0.5 µg/µl trypsin protease
0.5 µg/µl LysC protease

These buffers can be aliquoted and stored at -20°C.
*CAUTION: CAA and DTT are toxic. Prepare this solution in a fume hood and handle with gloves.*

**• Lysis buffer**

Prepare TFE lysis buffer containing 50% (v/v) 2,2,2-Trifluoroethanol (TFE), 300 mM Tris-HCl (pH 8.0).
*CRITICAL: This buffer should be prepared fresh. TFE is highly volatile, close caps directly after pipetting.*
*CAUTION: TFE is toxic. Prepare this solution in a fume hood and handle with gloves.*
*NOTE: The high Tris/HCl concentration (300 mM) in the lysis buffer assists in formaldehyde de-crosslinking* [2]
*NOTE: 50% TFE in the lysis buffer can be replaced by 50% acetonitrile for comparable results.*

**• Digestion buffer**

10% (v/v) TFE in ddH20. This buffer should be prepared fresh. Enzymes (trypsin and LysC) are added at a protein:enzyme ratio of 50:1.

**• SDB-RPS StageTip wash buffer 1**

1% (vol/vol) TFA in isopropanol. *CAUTION: TFA solutions are corrosive. Prepare the solutions in a fume hood and handle with gloves. This buffer is stable for >3 months at RT.*

**• SDB-RPS StageTip wash buffer 2**

0.2% (vol/vol) TFA. *CAUTION: TFA solutions are corrosive. Prepare the solutions in a fume hood and handle with gloves. This buffer is stable for >3 months at RT.*

**• SDB-RPS StageTip elution buffer**

1% ammonia, 80% ACN.

**• MS loading buffer**

0.2% TFA/2% (vol/vol) ACN. This buffer is stable for >6 months at RT.

#### Tissue preparation

Prior tissue collection via macroscopic dissection or laser-capture microdissection (LCM), all FFPE samples were deparaffinized and hematoxylin-eosine (H&E) stained as described below.

| **Deparaffinization** | | **H & E staining** | |
|---|---|---|---|
| **Step** | **Time** | **Step** | **Time** |
| Oven - 60 °C | 10min | Mayer's Hematoxylin | 1 min |
| Xylene | 10min | Water | 10 min |
| Xylene | 5 min | Eosin | 30 sec |
| 2 x 99 % ethanol | 5 min each | Water | 10 dips |
| 2 x 96 % ethanol | 5 min each | 70 % ethanol | 10 dips |
| 70% ethanol | 5 min | 96 % ethanol | 10 dips |
| Water | 10min | 96 % ethanol | 10 dips |
| | | 99 % ethanol | 10 dips |
| | | 99 % ethanol | 10 dips |

Areas of interest were determined via microscopic inspection by pathologists. For macro-dissected samples, an area of roughly 5 mm x 5 mm was collected by scraping with a razor blade (5 µm or 10 µm thick section). Note, tissue transfer into PCR tubes is enhanced with a few droplets of PBS on the scraped areas and subsequent pipetting. For LCM, FFPE tissues were mounted on PEN membrane slides to enable efficient cutting and collection. An area of approximately 1.5 mm x 1.5 mm (10 µm thick section, approx. 10,000 cells as calculated from dissected area × slide thickness / average mammalian cell volume of 2,250 µm$^3$, BNID 100434) was collected into adhesive caps and tissue transferred in lysis buffer into PCR tubes for direct in-solution protein extraction and digestion.

### PROTOCOL

#### *1. Tissue homogenization and formalin de-crosslinking (day 1, ~2.5h)*

- ☐ Add 100 µl lysis buffer (LB) to FFPE tissue collected into PCR tubes. Close PCR tubes with cap strips (see equipment).
  *CRITICAL: Make sure that caps are tightly closed.*
  *NOTE: We highly recommend working in PCR tubes. If samples were collected in different tubes or adhesive caps, transfer them into PCR tubes with lysis buffer. Make sure to collect all tissue pieces so that sample loss from transfer can be minimized.*
- ☐ Sonicate tissue (15 cycles in Bioruptor, high intensity, 30 s on/off cycle).
  *NOTE: Efficient sonication will lead to a homogeneous tissue powder. For some samples this will only be achieved in a second sonication step after 90 min boiling.*
- ☐ Centrifuge any condensation down
- ☐ Boil tissue at 90 °C for 90 min, no shaking.
  *CAUTION: TFE is toxic, work under fume hood.*
  *CAUTION: TFE is highly volatile and long heating times will result in overpressure during incubation. To minimize cap opening from overpressure during boiling, heat-resistant material (i.e. metal plate) can be placed on top. After boiling, a short 10 min cooling period to ~60 °C will reduce high pressure to safely remove samples from the heating block.*
  *NOTE: We found that overnight de-crosslinking at 65 °C offers a good alternative to avoid overpressure from high temperatures.*
  *NOTE: 50% TFE in the lysis buffer can be replaced by 50% acetonitrile for comparable results (see Fig. S1F-H).*
- ☐ Centrifuge any condensation down

#### *2. Protein reduction, alkylation and tryptic overnight digestion (day1/2, ~16h)*

- ☐ Add DTT (5 mM final) and incubate 20 min at RT, 1500 rpm.
  *CRITICAL: We noticed that during the long heating phase, buffer evaporation (if not avoided) can result in slightly different sample volumes after boiling. If necessary, adjust sample volumes with ddH20 to the original 100 µl and then add DTT.*
- ☐ Add CAA (25 mM final) and incubate 20 min at RT, 1500 rpm.
- ☐ Vacuum-dry to a remaining volume of about 20 µl completely (~45 min at 60 °C). *NOTE: Freeze or continue.*
- ☐ Add 80 µl freshly prepared digestion buffer including trypsin and LysC at an enzyme/protein ratio of 1:50. Incubate at 37°C 1500 rpm, overnight.
  *NOTE: Depending on tissue type and amount, enzyme amounts need to be determined empirically. We found that a tumor area of ~5 mm x 5 mm (10 µM thick section) resulted in 5-15 µg of total protein.*

#### *3. Peptide clean-up (day 2, ~1h)*

- ☐ Add TFA to a 1% final concentration to acidify the solution and inactivate trypsin and LysC. Mix by pipetting and spin down 5 min to pellet any debris.
  *NOTE: Freeze or continue.*
  *CRITICAL: A low pH is required for peptide clean-up by SDB-RPS StageTips (see below)*

- ☐ ***Peptide purification via StageTips***
- \- Prepare SDB-RPS StageTip with two layers:
- \- Load sample directly on SDB-RPS StageTips
- \- Wash with 200 µl wash buffer 1
- \- Wash with 200 µl wash buffer 2
- \- Change collection plate and add 50 µl elution buffer
- ☐ Vacuum-dry completely (~30 min at 45°C),
- ☐ Reconstitute peptides in 10 µl MS loading buffer. Measure A280 nm on a Nanodrop. Store peptides at -20°C until LC-MS analysis and inject 250-500 ng.
  *NOTE: A distinct 280 nm peak is indicative of a pure peptide sample.*


1 Rappsilber J, Mann M, Ishihama Y. Protocol for micro-purification, enrichment, pre-fractionation and storage of peptides for proteomics using StageTips. *Nat Protoc* 2007; **2**: 1896-1906

2 Kawashima Y, Kodera Y, Singh A, *et al.* Efficient extraction of proteins from formalin-fixed paraffin-embedded tissues requires higher concentration of tris ( hydroxymethyl ) aminomethane. *Clin Proteomics* 2014; **11**: 1-6
